# Perception Enhancement Using Visual Attributes in Sequence Motif Visualization

**DOI:** 10.1101/066928

**Authors:** Kok Weiying, Oon Yin Bee, Lee Nung Kion

## Abstract

Sequence logo is a well-accepted scientific method to visualize the conservation characteristics of biological sequence motifs. Previous studies found that using sequence logo graphical representation for scientific evidence reports or arguments could seriously cause biases and misinterpretation by users. This study investigates on the visual attributes performance of a sequence logo in helping users to perceive and interpret the information based on preattentive theories and Gestalt principles of perception. A survey was conducted to gather users’ opinion after being presented with several alternative design details to perform selected tasks on motif analysis. Analysis of results showed that there are improvements needed on the use of colour, negative space, size, and arrangement of the nucleotides, richness of information and interactivity aspect in a sequence logo visualization. These improvements can alleviate biases and misinterpretation of the results in sequence logo visualization.

## 1.0 Introduction

Visualization has become a very essential technique used by molecular biologists or bioinformaticians to present informative scientific findings. It helps in the exploratory analysis, the confirmatory analysis and the presentation of data [1]. Besides containing the design of techniques and tools for browsing, formulating and displaying the predicted outcomes or complex database queries, data visualization also contain the automated description and validation of the data analysis outcomes [2]. Information visualization can be viewed as a channel of communication from a dataset to the centre of cognitive processing. The power and usefulness of visualization in representing information of scientific data is largely due to the strength of human perception. Hence, besides functional consideration, human factor theories should be central to the design and development of visualization tools.

Gestalt theory proposed by the German psychologists Max Wertheimer, Kurt Koffka and Wolfgang Köhler in 1920s is one of the most widely used theories in perception. It explains how human tends to visually assemble individual visual element into groups or ‘unified wholes’ [3]. The use of visual attributes such as colours, symbols, shape, sizes and etc. are all dealing with how well the information can be represented in a piece of graphical representation. Thus, understanding the human visual system and perception mechanism are very important in improving the quality and quantity of information displayed in a graphical representation.

New visualization tools are continuously being proposed for various data visualization tasks. However, the lack of framework or defined theories in information visualization makes these tools difficult to validate and defend [4]. The greatest challenge most researchers faced while designing a visual tool is to decide on the graphical representation. They are concerned with the algorithms and how to transform data into a graphical representation especially in scientific data visualization. When it comes to deciding the best form of representation, models and principles on human factors are always being neglected and are mostly resort to their own aesthetic judgment.

Sequence logo is a visualization method introduced by Schneider and Stephens (1990) to visualize the conservation characteristics of the biological sequence motifs of DNA, RNA or protein [5]. Biological sequence motifs are short recurring sequence patterns that can be the transcription factor binding sites (TFBS) for DNA, the splice junction for RNA or the binding domains for protein molecules [6]. Identifying these sequence motifs are a big challenge for researchers as the sequence motifs are never exactly the same. Therefore, tools for discovering and visualizing sequence motifs are important for life scientist to solve various motif finding problems.

Previous studies have reported several critical limitations of using sequence logo for motif prediction tool evaluation and presenting motif analysis result. For instance, due to the lack of visual cues, it is difficult to obtain accurate interpretation of results from the visualization [7]. Therefore, heuristic rules are typically employed to make judgment or interpretation. Besides that, biases in decision making was found due to the limited information available on the graphical representation. Likewise, in another study, it was reported that using sequence logo to represent TF specificity might leads to misinterpretation of the prediction result [8]. For example, it was found that the appearance and information content of a motif shown as a sequence logo may not reflect its accuracy compare with the results obtained from wet-lab experiment. That can mislead users when performing comparison task. These limitations show that improvements are necessary in sequence logo graphical representation in order to support the interpretation of the results obtained.

The aim of this paper is to investigate the visual attributes performance in capturing user’s attention and perception while visualizing sequence logo based on the preattentive theories and Gestalt principles. Besides that, several modifications on the sequence logo are proposed to enhance the visual representation.

## 2.0 RELATED WORK

### 2.1 Sequence Logo

A sequence logo is a graphical representation that illustrates the specificity of sequences (i.e. motif) bind by a transcription factor protein. It is generated in two steps: (a) multiple-align the binding site sequences; (b) compute the relative frequency of nucleotides in each position of the multiple-alignment. The relative frequencies are subsequently used to compute nucleotides’ information content (in bits) using the Shannon’s information theory [5]. The information content values and relative frequencies of nucleotides in different positions are used to plot the sequence logo. In which the nucleotide letters in a position of a logo are sorted according to their relative frequency from the most frequent on the top to the least frequent at the bottom; and the total height of the four nucleotides letters (A, C, G, T) are determined by the information content value (conservation). Figure 1 shows an example of a DNA motif sequence logo generated by using WebLogo [9].

**Figure 1.**
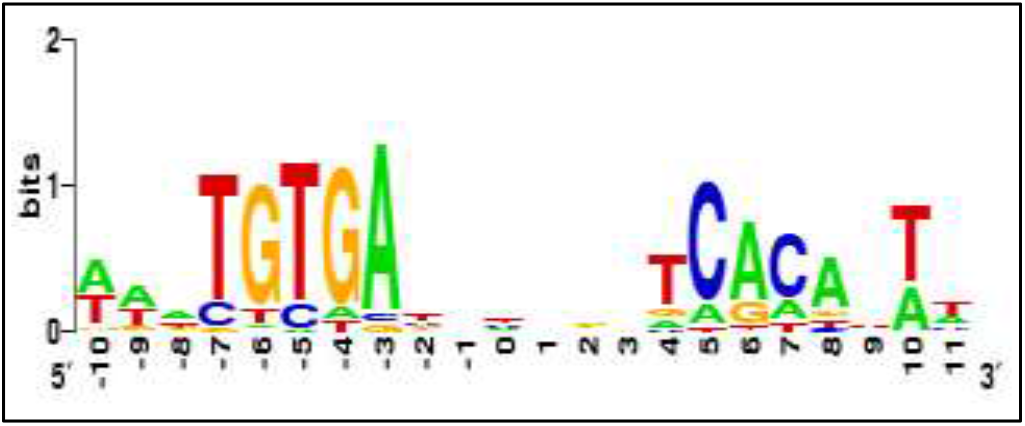
An example of a motif sequence logo. Shown is the Catabolite Activator Protein Binding Sites motif profile generated by WebLogo

From a sequence logo (Figure 1) we can obtain the following information: (a) the preferred binding site pattern bind by a transcription factor protein; (b) the order of predominance of the nucleotides at each position of the multiple-alignment; (c) the relative frequency of each nucleotide; (d) the amount of information content at each position of the multiple-alignment; and (e) the initiation point or cut point of the sequences. The core elements of a visualized DNA motif using a sequence logo are shown in Table 1.

**Table 1.** Core elements in a DNA motif Sequence Logo

| Element | Description |
| --- | --- |
| **Nucleotides** | The letters A, C, G and T which represents the four types of nucleotides (Adenine, Cytosine, Guanine and Thymine) found in DNA sequence. |
| **Colour** | Each nucleotide letters is associated with a distinct primary colour, monochrome or in grey scale. However, there is no standard with the colouring scheme used. |
| **Horizontal axis** | Shows the actual position relative to the genes where they are located or the position in the input multiple aligned sequences. |
| **Vertical axis** | Indicates the information content of a sequence position. The relative frequencies and conservation level of the nucleotides are shown in bits. |

(Source from: Lee & Oon, 2013)

Sequence logo is widely used as a method to present DNA motif analysis result such as: (a) to evaluate a newly proposed algorithm or tool for solving sequence motif discovery problem; (b) as a support evidence for the computational framework or the biological methodology; and (c) to visualize and compare the characteristics of the different binding site specification of a same transcription factor (TF) [7].

Weblogo is a web-based platform for sequence logo generation. It can generate the standard sequence logo for DNA, RNA, and protein sequences [9]. Several customized, specialized, or improved version of graphical representations basing on the original sequence logo have been developed to assist the visualization of specific motifs or functional sequence units [10]. Examples of the extension to the original sequence logo are such as: (1) CorreLogo which displays the correlation and potential base pairing in 3D graphical representation [11]; (2) RNA structure logo which combines the standard sequence logo with the mutual information of the base pairs [12]; (3) enoLOGOS which displays the standard sequence logo with the energy measurement, probability matrices and alignment matrices [13]; (4) BLogo which displays the statistical significant bias at each position in a sequence logo [14]. While many of these revised tools are useful, the original sequence logo remains the most popular. In this study, we employ the WebLogo tool to generate all sequence logos [10].

### 2.2 Visual Perception

Visualization process can be represented by a data encoding pipeline. The process of data encoding involves:

- Data transformation – converts raw data into a well-organized data format such as creation of new quantities or subset, modelling operations and etc.
- Visualization mapping – converts transform data into graphical representation with the use of suitable visual attributes
- Rendering – display the representation on screen

After the data is encoded and displayed, viewer will then interpret the graphical representation and this step is called the data decoding step which involves the perception and cognition process. Figure 2 shows the visualization process which involves the data encoding and decoding steps summarized by Wünsche [15]. Both visualization mapping and perception stages are connected by the visual attributes such as colour, shape, position and etc. A good visualization means the decoding of the information must be efficient and correct where maximum number of information should be perceived in a minimal time and relationship can be made between the data [15]. Thus, it is important to select the most suitable visual attributes to be used in the graphical representation as it will either help or hinder the user’s ability to interpret the data and facilitate the analysis tasks [16].

**Figure 2.**
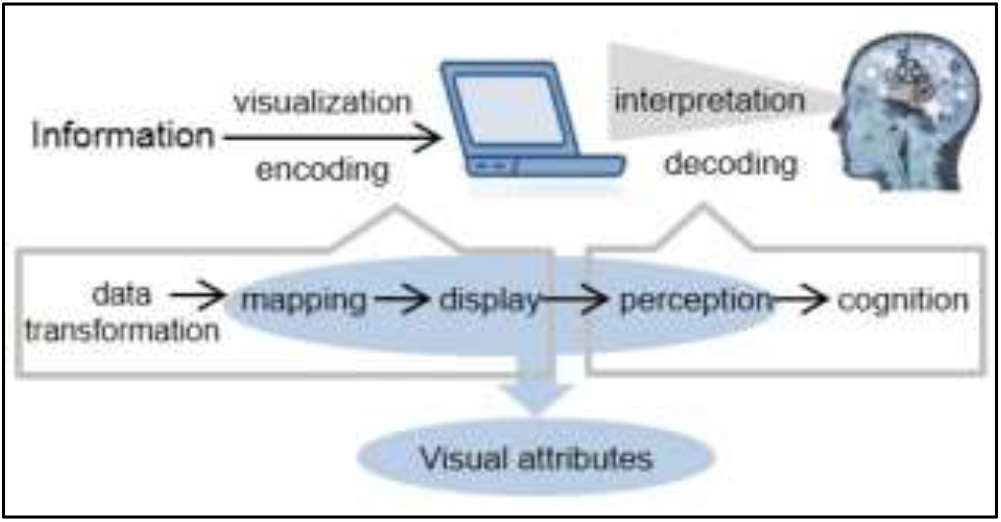
Visualization process

Visual attributes play an important role in capturing viewer’s attention and organizing or grouping visual display in the visualization process. Visual attributes such as orientation, length, closure, size, curvature, density, number, hue, luminance, intersections, terminators, 3D depth, flicker, direction of motion, velocity of motion and lighting direction are identified as preattentive visual features based on the five preattentive theory models [17, 18, 19, 20, 21]. According to the five well known models i.e. Feature Integration Theory, Textons Theory, Similarity Theory, Guided Search Theory and Boolean Maps Theory, low level visual processes can help to guide our attention in a larger scene and draw our eyes in order to have focus attention to a particular object in a scene [22]. Besides that, it can also minimize the mental capacity and help viewer to perceive the fundamental element presence in a graphical display effortlessly. Therefore, it is important to choose the proper visual attributes used in the graphical representation of sequence logo to draw viewer’s attention and correctly weight the perceptual strength of the data elements based on their encoded attribute values.

Moreover, low-level visual attributes are very useful in performing complex visual tasks such as the perception of depth, shape and gestalt whereas high-level visual attributes are involved in other higher order of tasks such as figure-ground and texture perception [15]. In Gestalt perception, the fundamental concept behind the principles is to explain how human tends to organize individual elements into groups [3]. This theory was initially used in the study of psychology but the concepts of the Gestalt theory have inspired many research areas such as image retrieval, visual design, graph drawing and etc. [23]. According to Koffka 1922, he stated that “the whole is greater than the sum of its parts” which shows that Gestalt is actually interplay between the parts and whole. The prominence on “wholes” had led the Gestalt psychologist to focus on defining the principles to explain about perceptual organization. In this study, we chose three of the principles that are related in the current design of sequence logo to test on the visual attributes performance.

- Law of similarity – human visual system tends to grouped elements which have similar attributes together in the same group. The use of colours and letters are used in sequence logo to represent the nucleotide Adenine, Cytosine, Guanine & Thymine;
- Law of figure and ground – the distinct element of focus will be seen as in front of the ground whereas ground means the background where figure placed will be seen as unformed material that seems to extend behind the figure. The use white colour background is used in the sequence logo to differentiate the nucleotide letters and the background of the logo;
- Law of proximity – objects that are near each other tend to be group together whereas objects that are further apart will be seen as less related or unrelated at all.

Thus, the performance of the visual attributes in guiding users to perceive the sequence logo based on these laws will be evaluated and some recommended improvements will also be tested to enhance the perception of the graphical representation.

## 3.0 METHODS

An online survey was conducted to determine the impact of introducing visual attributes in the current design of sequence logo. Purposive sampling was used in the survey in which international participants in the professions related to genetic, molecular biology, or bioinformatic were invited through email to participate in the survey.

The survey consists of three sections. Section A contains the basic demographic information, area of expertise, skill levels in using sequence logo and tools that have been used to generate sequence logo. Section B consists of 30 close-ended questions which are divided into 3 sub-sections: colour, layout and decision making. These questions would make performance comparisons related to the three visual attributes between the current and the modified design of sequence logo. Likert scale 1–7 is used to indicate participant’s opinion towards the comparisons. In addition to the questions related to visual attributes, participants were also being asked to compare and rate the perceived accuracy rate and quality level of visualized motifs predicted by two *de novo* computational tools i.e. MDSCAN [24] and MEME [25]. Section C consists of four open ended questions to gather participants feedback on the problems faced, satisfaction level and improvement needed on the current design of sequence logo.

## 4.0 RESULTS

A total of 52 participants from countries such as United States (48.1%), Malaysia (21.2%), United Kingdom (13.5%), Singapore (9.6%), Australia (5.8%) and Denmark (1.9%) participated voluntarily in the online survey. The participants involved were consist of researchers (46.2%), academician (21.2%), postgraduate students (30.7%) and bioinformatician (1.9%). From that 67.3% of the participants are male and 32.4% are female. Most of the participants are between 31 to 40 years old (44.2%); 34.6% are below 30 years old and 21.1% are 41 years old and above.

### 4.1 Colour of the nucleotide

One of the visual attributes that is commonly used for grouping and capturing viewer’s attention is the use of colour [27]. Colour helps viewer to discriminate objects that are differed from their surroundings and therefore is best for labelling and categorizing the information in a display. There are several factors to consider while choosing the right colour: the distinctness, unique hues, contrast with background, the number of colour used, field size, and colour blindness [26]. Choosing the right colour for data visualization will enhance and clarify the presentation whereas choosing the wrong colour will obscure, muddle and confuse viewer [27].

The default colours use by the Weblogo tool to identify the nucleotide letters are: A (green), T (red), G (yellow) and C (blue). We asked the participants to compare a logo with and without colours. From the survey, most of the participants agreed (46.2%) that the use of colours helped them identifying the nonconserved nucleotides in a sequence logo. 65.4% of the participants strongly agreed that it is easier to recognize nucleotides with colours compare with only black colour for all nucleotides. These results showed that the use of colour feature in a sequence logo is very important especially for preattentive processing and similarities grouping of the nucleotides. The use of black and white logo causes difficulties in perceiving the nonconserved nucleotides that are having small bits value. Figure 3 shows the comparison between coloured and black and white sequence logo.

**Figure 3.**
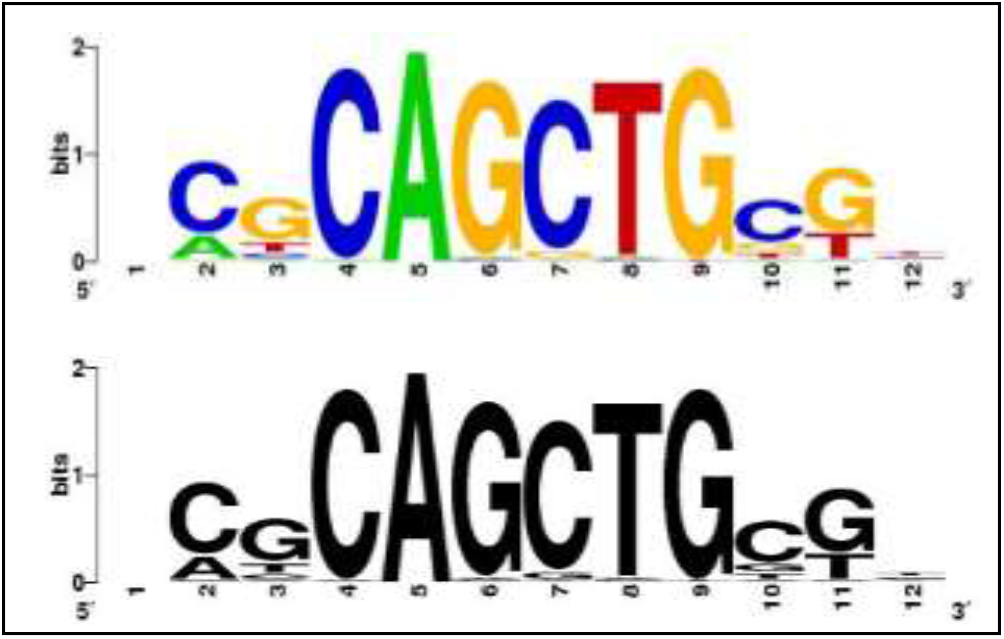
An example of coloured vs black and white sequence logo

We also determine whether the use of only one colour to represent the most conserved nucleotide at each position of a sequence logo would improve users’ task. The results from the paired sample t-test shows that there is a significant difference in the ease of identifying conserved nucleotides by using only one colour for the most conserved nucleotide compare to original design of sequence logo (t=-7.005, p<0.001). The use of a single colour to represent the consensus sequence guides user in the preattentive processing while visualizing the sequence logo. Besides that, it will also assist in the grouping of the most conserved nucleotide and the non-conserved nucleotide based on the law of similarity in Gestalt perception. Figure 4 shows the comparison between the original design with the proposed modification.

**Figure 4.**
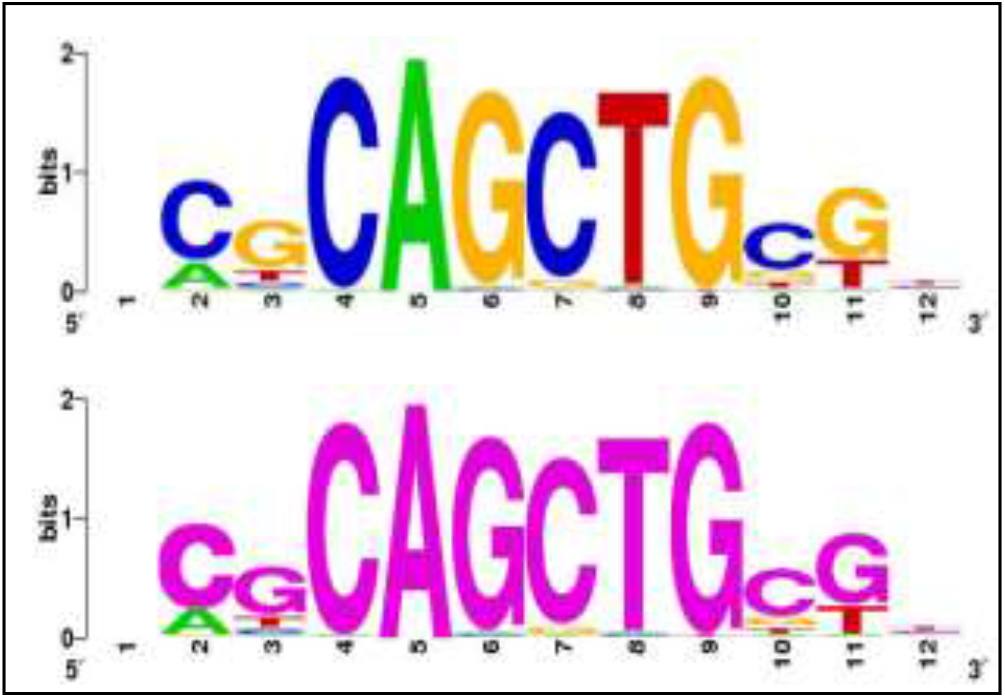
Comparison between the use of different colours and same colour for the most conserved nucleotide in sequence logo

We compared on the use of bolded letters versus the use of mono colour to represent the most conserved nucleotide letter in each position of a sequence logo. The result shows a significant difference in the ease of identifying the conserved nucleotide by using same colour compare with the use of bolded letters (t=3.525, p=0.001). Same coloured consensus sequence produce a “pop out” effect in the graphical representation which easily discriminate the conserved and non-conserved nucleotide compare with the use of bolded letters. This will guide viewers to rapidly and accurately identify the conserved nucleotide shown in a sequence logo (see Figure 5 as an example).

**Figure 5.**
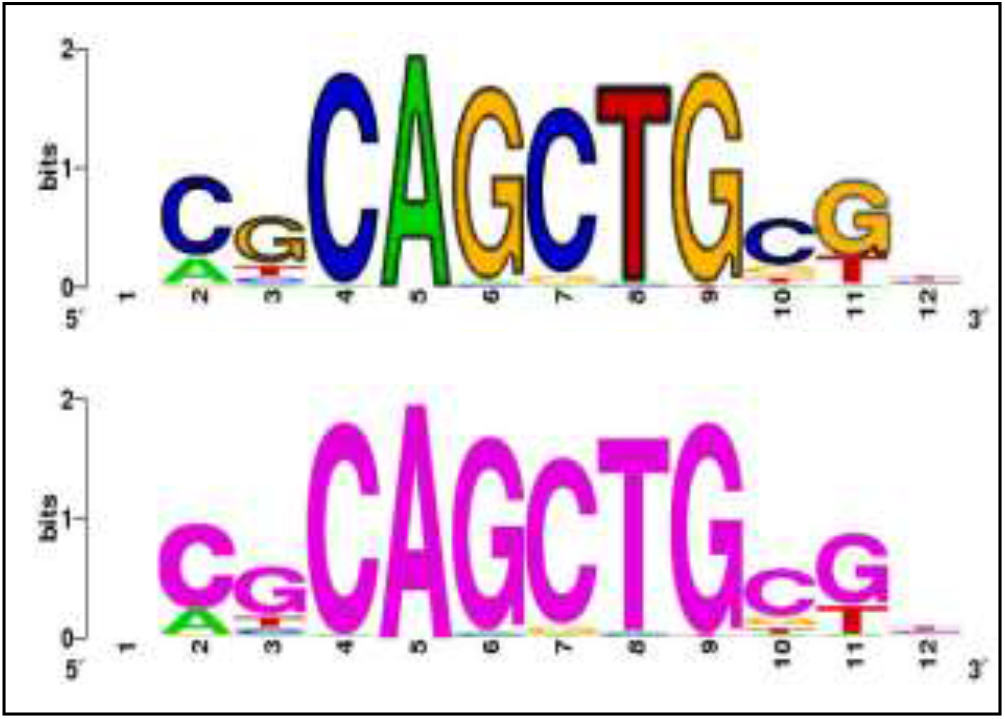
Comparison between the use of bolded letters and mono-colour to represent the most conserved nucleotide in each position of a motif.

Hence, improvement in the use of colour in indicating the conserved or non-conserved nucleotide is needed as it will act as a feature to capture user’s attention and helps to distinguish different categories of the data presented in a sequence logo.

### 4.2 Negative space in the Sequence Logo

Negative space is the empty space which usually serves as the background in a graphical display. According to the law of figure and ground, human visual system has the perceptual tendency to separate the object (element of focus) from the background [3]. Thus, graphical design with well-composed negative space will help viewer to identify the important information or object in a display effortlessly. The proper use of negative space will help viewer to focus their attention to the object and pay less attention to the background. The more contrast between the figure and ground, the easier viewer can distinguish the objects. Moreover, negative space can also serve as the grouping element in a display. According to the law of proximity, the relative spacing between the columns or rows will dramatically affect our grouping ability [3]. Hence, the width of negative space will affect our ability to group the objects vertically or horizontally.

In the questionnaire, participants were asked on the use of spaces between neighbouring stacks of nucleotide letters. Most of the participants slightly disagreed (36.5%) that the spacing between the stacks in the design of sequence logo are too near which causes it to look messy. Besides, the participants perceived no significant difference between the sequence logo without spaces (M=3.88, SD=1.517) and the modified design with spaces (M=4.06, SD=1.474) between the nucleotides stacks (see Figure 6 as an example). This perhaps is due to the use of white background colour in the current sequence logo design which provided a good contrast between the stacks and the coloured nucleotides for users to perceive the important information needed.

**Figure 6.**
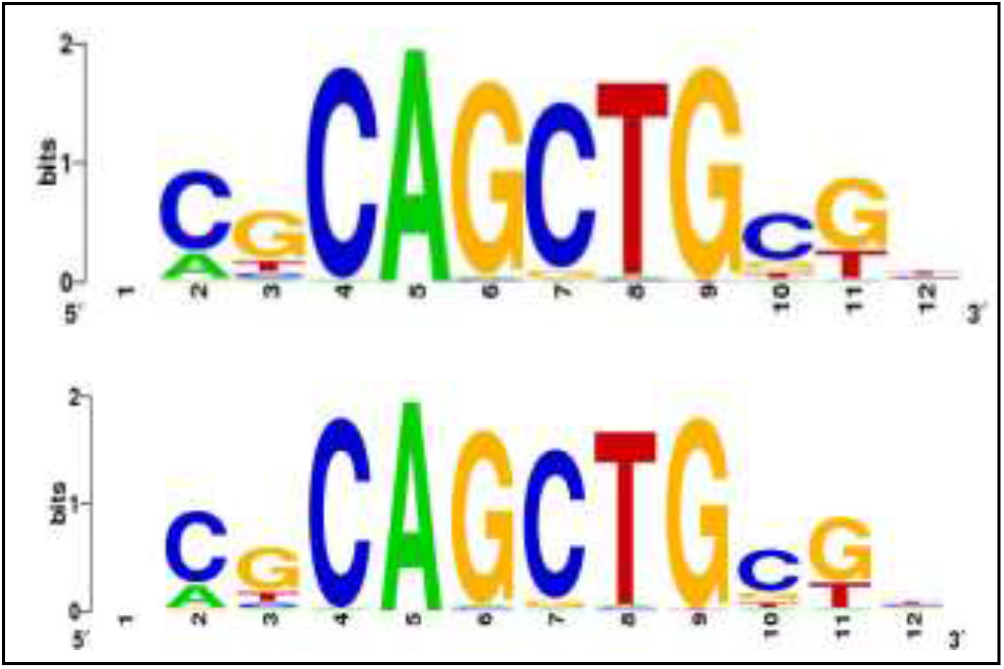
Comparison between the sequence logo without spaces and spaces between the nucleotides stacks

We also proposed to place a thin line between neighbouring stacks of nucleotides to create the effect of grouping the stacks. The result shows that there is a significant difference in the use of empty spaces in contrast to the use of thin lines (t=3.578, p=0.001) (see Figure 7 as an example). Based on the law of proximity, the existent of white space between each position helps to easily group the stack of nucleotide letters together whereas the use of thin lines may cause confusion to the viewer especially if it involves a long sequence motif.

**Figure 7.**
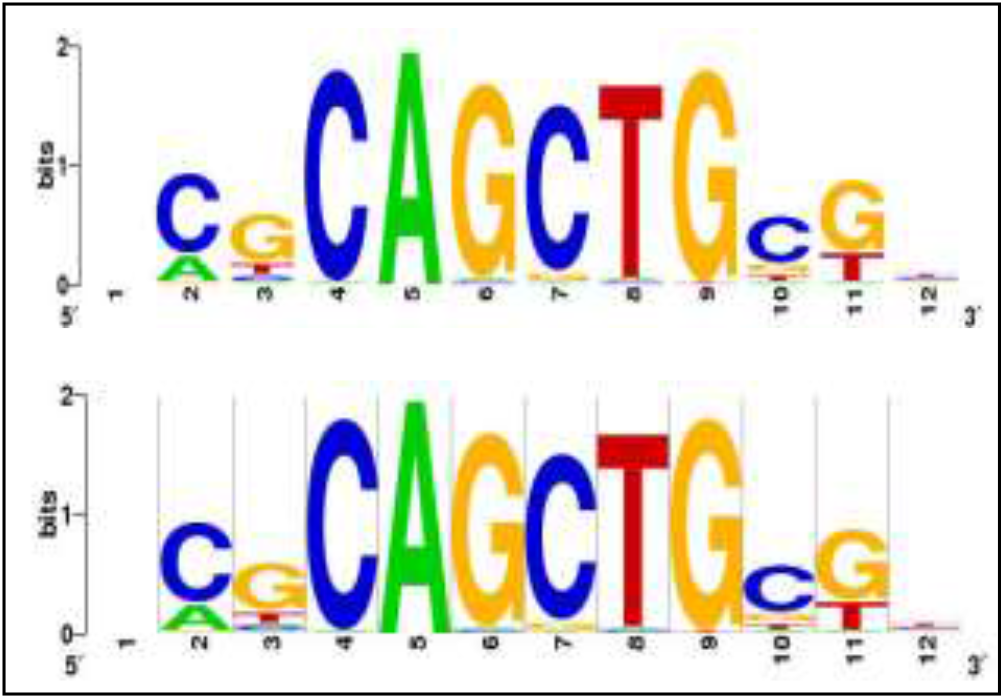
Comparison between the sequence logo with spaces and thin lines in between the stacks of nucleotides

The presence of negative space in a sequence logo can help viewer to easily identify the stack of nucleotides at different position. Therefore, it is not recommended to place a thin line between each stack of nucleotide in the sequence logo to avoid unwanted confusion to the viewers.

### 4.3 Size and arrangement of the nucleotide

The total height of the four stacked nucleotide letters and their vertical ordering in the stack represent the total information content and nucleotide’s relative frequency at each position, respectively. Most of the participants agreed (73.1%) that it is difficult to determine the conservation of different nucleotides at the same position when their relative frequencies are almost identical. But only 25% of the participants slightly agreed to the idea of displaying nucleotides’ relative frequency values in a sequence logo.

Next we investigate the possible weaknesses of using “size” of nucleotide letters for comparison purposes. In the questionnaire, participants were asked to compare two sequence motifs predicted by MDSCAN and MEME with an annotated motif and rate the accuracy and the quality of the motif based on their perception. MDSCAN has a better prediction performance than MEME due to the higher f-measure [7]. The result shows that there is a significant difference in the perception of the accuracy level (t=4.592, p<0.001) and quality level (t=2.704, p<0.01) of the two predicted motifs compare with the annotated ones.

Figure 8 shows the comparison between the three sequence logos generated: (a) annotated motif; (b) MEME; (c) MDSCAN.

**Figure 8.**
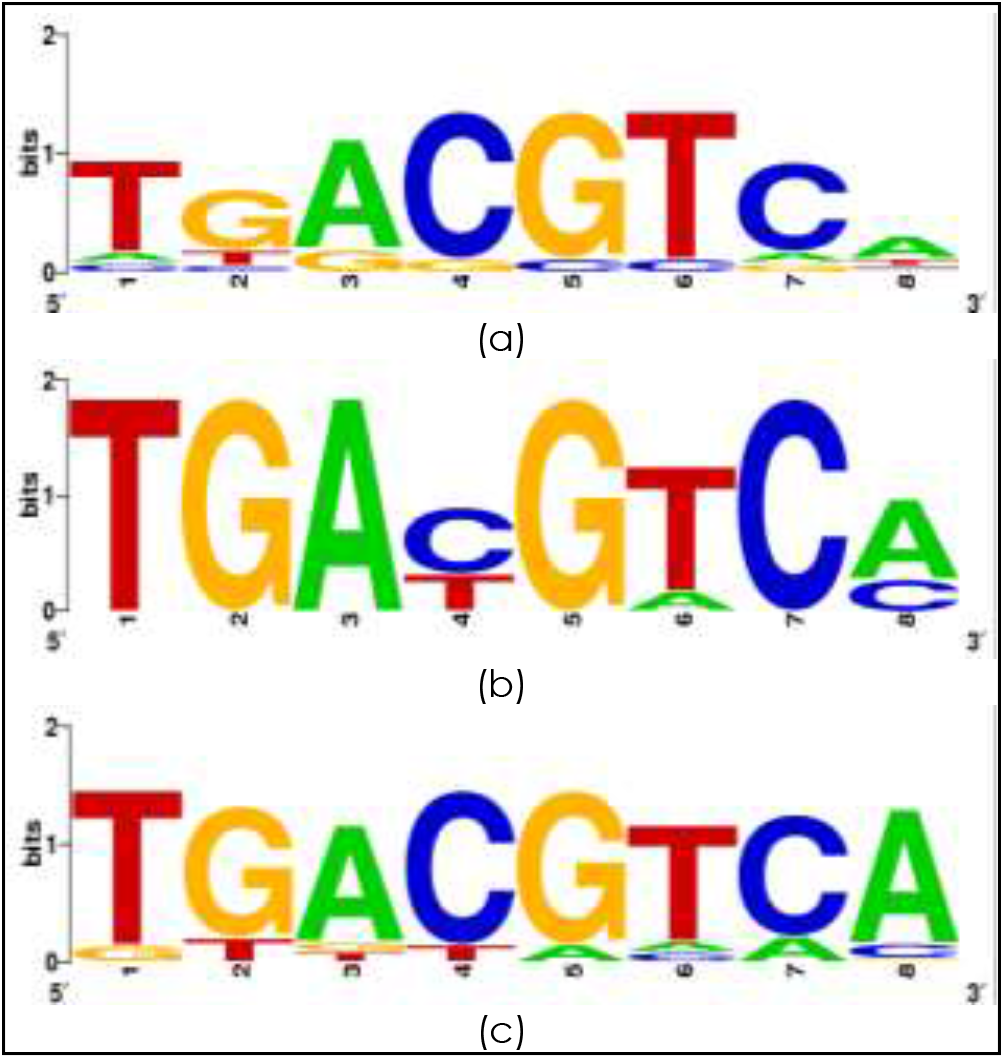
Comparison between the sequence logo generated by two de novo computational tools (b) MEME; (c) MDSCAN with the (a) annotated motif

This result is expected as the annotated motif is marked resemblance to the motif predicted by MDSCAN even though the motif obtained by MEME has many tall conserved nucleotides (high information content). Previous evaluation study also reported a similar issue of using the height of the stacked nucleotides for decision making and the lack of information for accurate interpretation of result [8]. This study has further support the finding of our previous findings where the current design of sequence logo is not accurate when is used to evaluate the predictive ability of computational tools and have to rely on heuristics rules to interpret result obtained [28].

We propose two modifications as a mean to improve the accuracy of reading information content values in a sequence logo. The first is by introducing horizontal grid lines and the second is to display a nucleotide’s bit value when the mouse cursor is hover over it (shown in Figure 9). We found statistical significant difference in participants’ preference in which they preferred the second over the first method (t= 3.796, p< 0.001). The present of bits value can help to clearly indicate the relative frequency of the nucleotides at each position and avoid misinterpretation of the results by the viewer.

**Figure 9.**
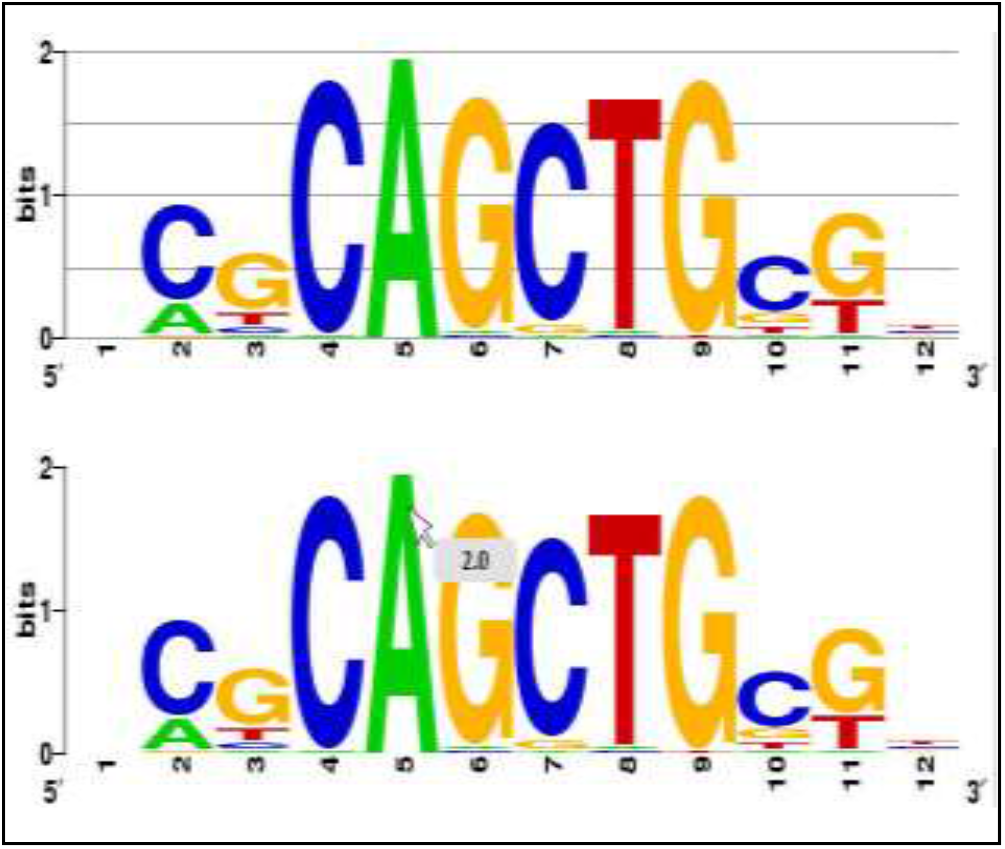
Comparison between the horizontal grid lines with mouse hover function to identify the bits value of the nucleotides

Other than the size, some problems were found in the arrangement of nucleotides at each position. It is known that the most dominant nucleotide is arranged at the top most position of a stack of nucleotide [5]. However, it is found that if only two nucleotides are presence at a position with the same relative frequency, the arrangement of the nucleotides is based on alphabetical order. We asked participants if two or more nucleotides with the same height at the same position would cause difficulty in decision making. Most of the participants agreed (55.8%) that this arrangement will mislead them especially when no extra information is shown in the sequence logo to indicate that both of the nucleotides have identical relative frequency. Moreover, majority of the participants agreed (46.2%) that it will be confusing, especially for novice users, when a position in a sequence logo is left blanked whenever the nucleotides have the same relative frequency (i.e. zero information content value).

Our findings showed that although the size and arrangement can help to identify the relative frequency and conservation of nucleotides in a sequence logo, there are some limitations in the graphical representation which can cause misinterpretation of the results obtained. Thus, it is necessary to improve the visual attributes of the current sequence logo design.

## 5.0 CONCLUSION

This study has identified some limitations on the visual attributes performance found in the current sequence logo design. Although the use of colour can help to categorize the nucleotides but confusion may occur if the colour of the nucleotides are the same. The negative space, size, and arrangement attributes in the current graphical representation help in preattentive processing and grouping of the information. However there are still some limitations with the use of these visual attributes. A more detailed feature comparison is needed to investigate the most suitable colours to represent the conserved and non-conserved nucleotides. In addition, further study is needed to determine the most suitable graphical representation which can clearly indicate nucleotides’ relative frequency.

Our findings also found that it is necessary to enrich the amount of information display on a sequence logo and provide interactivity feature which can assist both novice and expert users in the interpretation of a sequence logo. These improvements would alleviate the problems of biases and misinterpretation when using sequence logo for various motif analysis related tasks.

## Acknowledgement

This study is supported by the Ministry of Higher Education Research Acculturation Grant Scheme RAGS/b(4)/926/2012(27). We would like to thank all the participants involved in this study for their valuable time and opinion.

